# Evolution of a confluent gut epithelium under cyclic stretching

**DOI:** 10.1101/2021.07.10.451888

**Authors:** Lauriane Gérémie, Efe Ilker, Moencopi Bernheim-Dennery, Charles Cavaniol, Jean-Louis Viovy, Danijela Matic Vignjevic, Jean-François Joanny, Stéphanie Descroix

## Abstract

The progress of food in the gastrointestinal (GI) tract is driven by a peristaltic motion generated by the muscle belt surrounding the GI tract. In turn, the response of the intestinal epithelial cells to the peristaltic stresses affects the dynamics of the epithelial structure. In this work, we study the effect of cyclic stretching (0.125 Hz, 10% strain) on the spatial organization of the intestinal epithelium using intestinal cells deposited on a flat elastomeric substrate to mimic the peristaltic motion *in vitro*. A confluent monolayer of Caco-2 cells is grown on a PDMS chip to probe the morphological and orientational response of the tissue to cyclic stretching. The PDMS chips are either covalently or non-covalently coated with laminin to recapitulate the basement membrane. We observe a significant orientational response where the cells rearrange their long axes perpendicular to the stretching direction for both coating conditions. The experiment is modeled by a vertex model where the cells store elastic energy with varying strain and effectively have a rotational diffusive motion through rearrangements of their shapes. The model also predicts a transition between the perpendicular orientation and orientation at an oblique angle determined by the level of the cell elastic anisotropy. It provides a general framework to study cell response and relaxation dynamics under cyclic stretching across different cell types. We also discuss potential relevance of peristalsis in determining planar cell polarity in 3D architectures.

---

The small intestine is a major organ of the human body being at the same time the place of the nutrient absorption and a barrier against external entities. These functions are in part achieved by the epithelial monolayer that faces the intestinal lumen. The epithelium is attached to the basement membrane, composed of laminin and collagen IV. The basement membrane offers structural support for the epithelium and separates it from the stroma. The stroma consists of cellular components such as fibroblasts, immune and endothelial cells, and a supporting network of extracellular matrix (ECM).

The intestine is a complex organ with unique 3-dimensional structure and dynamics. It possesses an intricate topography: composed of finger-like protrusions, called villi, and crypts that descend into the stroma. These different regions are correlated with a well-defined patterning of various cell types: the intestinal stem cells are confined at the bottom of the crypts, whereas most of the differentiated cells migrate actively from the crypts to the tip of the villi where they are extruded from the monolayer [1]. In addition to being regulated by a wide range of topographical cues, the small intestine epithelium experiences various mechanical forces in both physiological and patho-physiological states [2]. In particular, intestinal epithelial cells are exposed to shear stress, cyclic deformation, and strain associated with villi motility. These forces affect cell properties, such as proliferation or adhesion, and tissue properties, such as the intestinal barrier function [3–5]. A significant source of mechanical stress is the peristaltic motion that arises from two layers of smooth muscles, the outer layer being oriented longitudinally and the inner one is oriented circumferentially [6]. Their contractions take place at different frequencies and amplitudes depending on the food uptake and are highly coordinated to act as propulsive forces and to mix the lumen content [7].

Over the last decade, there have been remarkable efforts to recapitulate *in vitro* complexity of gut anatomy. The most commonly used model is organoids generated from mouse intestine. They spontaneously form protrusions resembling crypts and exhibit normal cell differentiation and localization of intestinal stem cells. However, the lack of villus domain and organoids’ closed structure are clear limitations [8, 9]. Organ-on-chip technology is an appealing alternative to organoids, as it aims at recapitulating multicellular architectures, physical, biochemical and geometrical features of a specific organ on a microfluidic chip [10, 11]. Two main approaches are undertaken to develop a relevant gut-on-chip model. The first approach is to build a 3D scaffold that replicates the villi/crypt structure on which cells are seeded [12–15]. On such 3D scaffolds, cell segregation along the crypt-villus axis is achieved either by implementing growth factor gradients [16] or with the addition of stromal cells [17]. The second approach relies on applying external forces on cells to mimic the intestinal dynamics, which has been studied in 2D systems only [18–21]. In most models, cells are grown on deformable membranes, and cells are subjected to both stretch and shear stresses [3, 22]. After about 100 hours of stretching, cells spontaneously form folds resembling villi, and epithelial barrier function gets improved compared to no strain conditions [23]. The transcriptomic study showed that cells grown on those chips resemble the *in vivo* conditions more closely than organoids [24].

These recent papers have highlighted how mechanical forces applied to cells are instrumental to develop a relevant *in vitro* microphysiological gut model. While most studies address how cyclic stretching that mimics peristaltic motion impacts intestinal cell proliferation or differentiation, it remains unclear how the tissue organization is affected by these forces. Here we focus on the effect of peristaltic motion on the intestinal epithelium. To this aim, we developed a novel stretching device compatible with microfluidic chips that recapitulates both the basement membrane and the intestinal epithelium. We study two coating conditions of the basement membrane: laminin is either covalently or non-covalently coated on the PDMS chip. Next, we apply cyclic stretching on the chip where the intestinal epithelium is subject to a 10% strain at 0.125 Hz mimicking the *in vivo* peristaltic motion [21]. We next characterize the monolayer response to cyclic stretching in terms of cell spatial organization and morphology within the tissue for a wide range of stretching times to explore the kinetics of the tissue response. The most significant response of the tissue appears in the cell reorientation: a fraction of the cells orients their long axis perpendicular to the stretching direction. We model this behavior using a vertex model and combined it with an overdamped dynamics of the cell reorientation. Accordingly, we identify the effects of cell anisotropy and activity in determining cell orientation under peristalsis-like stimuli.

## Results and discussion

### Cell Monolayer Culture and Stretching Device

To assess the influence of stretching and cyclic stretching on the intestinal epithelium, we developed a customized stretching device along with dedicated microfluidic chip fabrication (Fig. 1A). The stretching device is compatible with conventional cell culture conditions and can accommodate 3 chips in parallel, and its mechanical strain and frequency can be tuned at will. In comparison, the most commonly used model of dynamic gut-on-chip is based on the use of a central cell culture chamber separated by a thin porous membrane flanked with two vacuum lateral channels that are used to perform cyclic stretching [25]. Those devices have shown their potential for different cell types [23, 26] as they allow an efficient strain application. However, they require complex microfabrication and microfluidic-based fluid handling. Here, we develop an original microfabrication approach to design open chips similar to a conventional cell culture dish compatible with stretching. The bottom of the chip where the cells are grown could be either flat as in the current work or 3D structured. These chips are made of polydimethylsiloxane (PDMS), a common cytocompatible elastomer used here as cell culture substrate (Fig. 1B). The PDMS prepolymer and curing agent ratio is set to 10 : 1 (w/w) with a Young modulus around 1.63 MPa [27]. Stretching creates a biaxial strain at the center of the chip with a longitudinal elongational strain 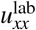, and a transverse contractile strain 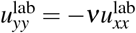 where the measured Poisson ratio of the substrate is *ν* = 0.4. We quantify the cells response to cyclic stretching only at the central part of the chip where the deformation can be considered homogeneous (Fig. 1C).

**Figure 1.**
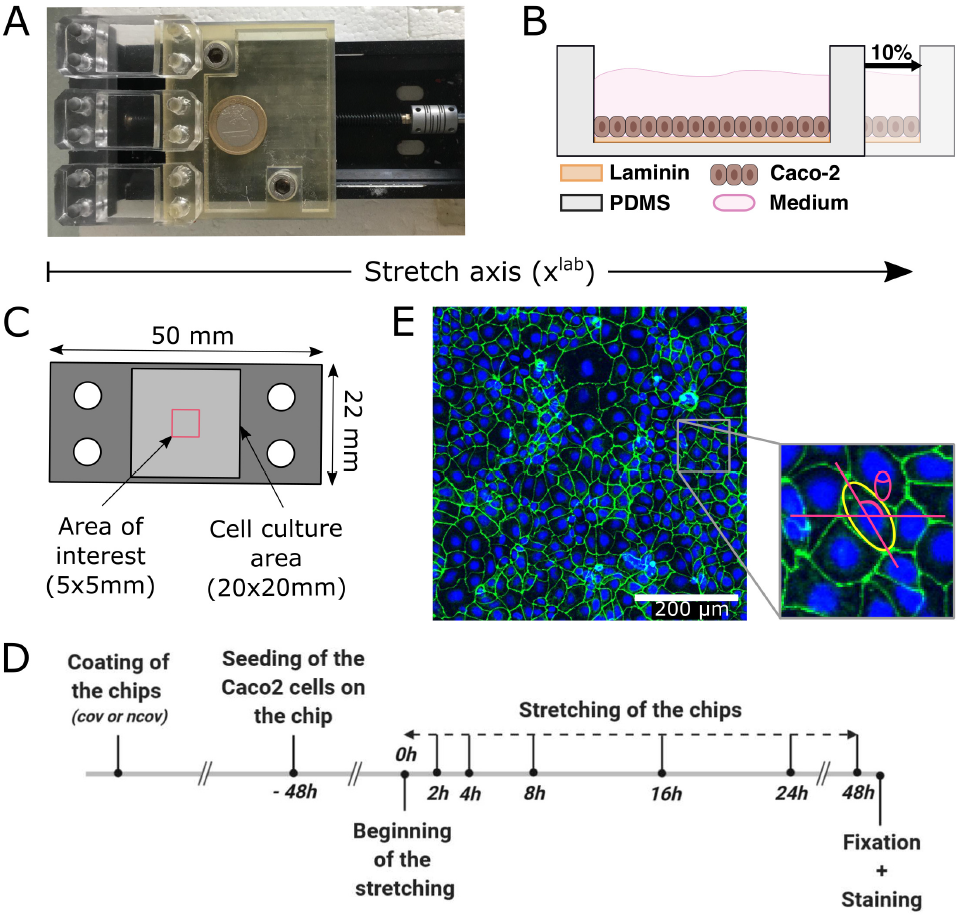
(A) Picture of the homemade stretching device allowing the stretching of three chips in parallel. (B) Schematics cross section of the chip representing the BM, the cells as well as the stretching deformation. (C) Schematics of the chip representing the area of interest where imaging is performed. (D) Timeline of the experimental workflow: from the chip coating to cell staining. (E) Picture of the confluent Caco-2 monolayer after 24 hours of stretching with a laminin covalent coating, using a 10x confocal microscope. The tight junctions are stained using ZO1 antibody (in green) and nucleus staining is performed using Hoecht (in blue. The insert shows the ellipse fit by ImageJ to estimate the cell long and short axis as well as cell orientation angle (*θ*) with respect to the stretching direction.

To reproduce the composition of the intestinal mucosa, we first recapitulated the intestinal basement membrane (BM) by coating the chips with laminin (Fig. 1B), the predominant BM glycoprotein [28]. The cell response to ECM deformation is mediated by focal adhesions (FAs) that link the cell to the substrate. Focal adhesion protein, paxillin, is activated during cyclic deformation of intestinal cells [29]. To model *in vitro* the cell-BM interactions, PDMS could be coated with adsorbed BM proteins such as laminin or collagen IV. More recently, Wipff et al proposed an original approach to covalently coat ECM proteins to stretchable PDMS membrane, which circumvents some limitations of non-covalent coating in particular its poor stability [30].

Here, we tested both coating procedures to assess how the coating conditions affect the epithelium response to cyclic stretching, assuming that a more stable attachment of laminin to the surface could improve cell adhesion. More precisely, the PDMS coating is performed either covalently, through PDMS silanisation coupled by glutaraldehyde crosslinking (as described in [30]), or non-covalently, through physisorption after PDMS activation by plasma treatment. To recapitulate the human intestinal epithelium, we grew the human colon carcinoma cell line (Caco-2) 48 hours on those chips so that they reach confluence before stretching (Fig. 1D). The Caco-2 cell line has been selected because it generates a tight epithelial monolayer and it is a common model to study the intestinal epithelium [3, 15, 31, 32].

### Effect of uniaxial cyclic stretching on epithelium morphology

To asses how stretching affects epithelial cells, we subjected the epithelial intestinal monolayer to a cyclic stretching, mimicking the intestinal peristalsis *in vivo*. We implemented a sinusoidal strain 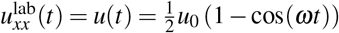 and 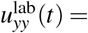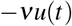 with angular frequency *ω* = 0.125 Hz ·2*π* and magnitude *u*_0_ = 0.1, imposed during a period ranging from 2 to 48 hours to measure the kinetics of the intestinal epithelium response. As a control, we used similar chips seeded with the Caco-2 cells but left deformation-free during the experiment.

In order to probe morphological changes in the tissue, we stained tight junctions using ZO1 (Zonula occludens-1). In both conditions, Caco-2 cells formed a tight monolayer with prominent tight junctions between cells. This epithelial monolayer was tightly packed with columnar cells with a hexagonal-like shape at their apical side characteristic of in vivo intestinal epithelium (Fig. 1E). We analyzed cell circularity, perimeter, and area using Tissue Analyzer plugins in Fiji software. Cells were fitted by an ellipse from which we measured their long and short axes to infer their aspect ratio and the orientation of their long axis with respect to the stretch direction.

We first examine if the laminin coating strategy affects intestinal cell morphology in the absence of stretching. Indeed, it has been shown that laminin surface density can be increased by covalent coating [33] compared to non-covalent coating, which could consequently affect cells spreading. We observed that the cell area was similar on covalently and non-covalently coated PDMS even after 24 hours of culture. This result suggests that the different coating strategies do not affect cell spreading. We next examined cell response to cyclic stretching. We found that stretching does not affect the morphology of the intestinal cells, even after a long stretching period. In particular, the mean area, perimeter, aspect ratio, or circularity of cells are similar between stretched and control conditions, as well as between covalently and non-covalently bound laminin (Fig. 2 A-D). Similarly, we did not observe any change of the monolayer height between different conditions (data not shown).

**Figure 2.**
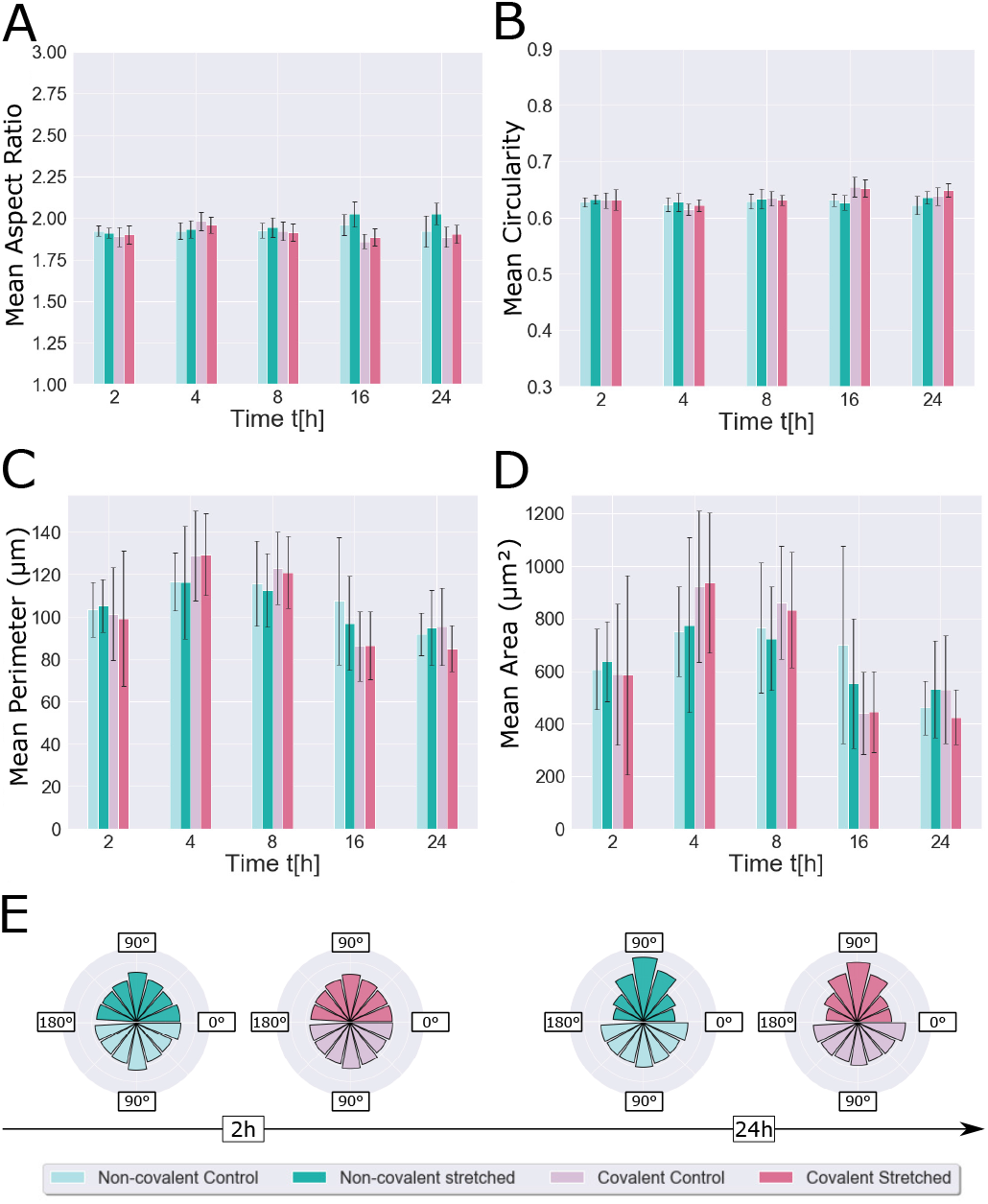
Variation of shape parameters of epithelial cells for different analysis times *t* under cyclic stretching with 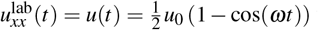 with *ω* = 0.125 Hz ·2*π* and magnitude *u*_0_ = 0.1. The mean aspect ratio (A), circularity (B), perimeter (C) and area (D) are plotted as a function of time for covalent and non-covalent coating and stretched and control conditions. These data have been collected from 15 images (5 images per chip) which corresponds approximately to 20,000 cells analyzed per condition. We calculated the mean parameter per image, and the error bars are obtained by the standard deviation between images of the same condition. All cell shape parameters remain constant over time and do not vary as functions of the stretching conditions. Figure (E) shows the angular distribution of cell orientation after 2h (left) and 24h (right) of stretching for covalent and non-covalent coating and stretched and control conditions. The percentage of cells perpendicular to the stretching direction increases with stretching time.

Nevertheless, after 48 hours of cyclic stretching, patches of cells start to detach from the cell substrate that is non–covalently coated with laminin, while no detachment is observed with covalently-coated laminin. Similar observations have been reported for human aortic smooth muscle cells grown on sub-millimeter hemi-channels made of glass which is either covalently or non-covalently coated with collagen I [34]. Higher substrate curvature or cell contractility results in increased cell detachment, which is likely due to an imbalance between cell-generated forces and the strength of cell-cell and cell-substrate adhesions. As our substrates are flat, cell detachment is likely due to an imbalance between stretch-induced cell forces and cell-substrate adhesion strength. These results are in good agreement with the lower stability of the non-covalent coating reported elsewhere [33].

Altogether, our results show that the cell shape parameters are insensitive to a cyclic stretch mimicking *in vivo* peristalsis regardless of the laminin coating conditions and remain constant over time. However, we noticed an anisotropic response of the cells measured by the orientation angle of their long axis, which we examine next in detail.

### Epithelial cells reorient perpendicular to principal strain direction under cyclic stretching

During periodic stretching, the cells are under varying strain such that there is no unique rest position on the stretching path from the reference state (non-stretched, 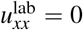) to the stretched state. Thus, the cells continuously feel the varying mechanical stimuli, and store elastic energy. Our results show that, in these conditions, a significant fraction of cells orient perpendicular to the stretching direction. As illustrated in Fig.2 E, after 24 hours of cyclic stretching, ∼20% of the total cell population orients between 80-100° to the stretching direction.

Previous works report a passive response to a finite non-varying stretch in which cells tend to orient parallel to the stretching direction [35, 36]. We also performed static stretch experiments that are detailed in the Supplementary Information (SI) Section 2 and observed a similar passive response. However, if the chip remains under constant strain *u*(*t*) = *u*_0_ during several hours, we observe a dissipative relaxation of cells orientation without any stored elastic energy, showing the reversibility of orientational behavior. Starting from an orientation that was parallel with the stretching direction, the ensemble of cells recovers orientational isotropy, with an orientational relaxation time *τ*_rel_ ∼ 5 − 7 hours. In contrast, during cyclic stretching, the cells can store elastic energy, as discussed in the following.

To model the orientational dynamics under cyclic stretching, we make the following assumptions: (i) Cells deform together with the substrate and hence passively follow the stretch direction at short times (*t* ∼ *τ*_str_) where *τ*_str_ = 2*π/ω* is the period of stretching. (ii) At long timescales (*t* ≫ *τ*_str_), the cells undergo a rotational diffusion. In the absence of strain, this leads to an isotropic distribution of orientations. As we exert cyclic stretching, the cells are subject to an angle-dependent potential as detailed below.

We write the total elastic energy stored in the tissue under varying strain using a vertex model. For a tissue containing *N* confluent cells, the total mechanical energy is given by *E*_tot_ = *E*_tot_({**r***_i_*}) [37–39],

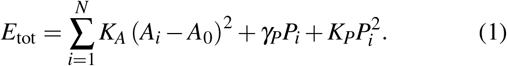

Each cell *i* at position **r***_i_* has a preferred area *A*_*i*,0_ = *A*_0_ that we consider identical for all cells. If a cell has an area *A_i_* different from *A*_0_, there is an energy cost with a positive modulus *K_A_*. The second term is a line energy proportional to the cell perimeter *P_i_*. At linear order, the line tension is *γ_P_P_i_*. This line tension includes contributions both from cell-cell adhesion and individual cell cortical tension. Because of the cell activity, the line tension depends on the cell perimeter and this leads to a correction 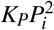. For stability reasons, the perimeter modulus *K_P_* is also positive. The cell has therefore a preferred perimeter *P*_0_ = −2*K_P_/γ_P_*. Using assumption (i) that the cells follow the deformation of the substrate during one cycle of stretching, we directly express the strain in the reference frame of the cells. Taking the long-axis and the short-axis of the cells respectively along the *x* and *y* directions (Fig. S1), we obtain *u*_*xx*_(*t*) = *u*(*t*) (cos^2^ *θ* − *ν* sin^2^ *θ*), *u*_*yy*_(*t*) = *u*(*t*) (sin^2^ *θ* − *ν* cos^2^ *θ* and *u*_*xy*_(*t*) = *u*(*t*)(1 + *ν*) cos *θ* sin *θ*. As a result, the area of the cell *A_i_* varies as *A_i_* → *A_i_*(1 + *u*(*t*))(1 − *νu*(*t*)) and it is independent of the orientation. By contrast, the amount of perimeter deformation depends on the orientation of the cells. To obtain the energy change per cell upon stretching, we make the meanfield approximation that all shape properties of the cells except their angular orientation can be treated as identical. This is supported by the fact that we do not observe a significant correlation between the other shape parameters and the orientation angle in all experimental conditions. As a result, we write the energy stored per cell as:

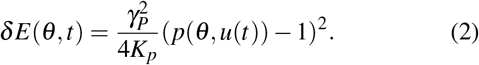

where *p_i_*(*θ*_*i*_, *u*(*t*)) = *P*_*i*_(*θ*_*i*_, *u*(*t*))/*P*_0_ is the ratio of the perimeter of *i*^th^ cell with orientation *θ_i_* under strain *u*(*t*) with respect to the preferred perimeter value *P*_0_ under no strain (see SI Section 1 for details).

As we observe in static stretching experiments (where *δ E*(*θ, t*) = 0), the rotational diffusion is a much slower process than the stretching frequency, i.e., *D_b_* ≪ π^2^*ω* where *D_b_* is the basement membrane dependent rotational diffusion constant, and hence the mechanical energy gain can be averaged over one period of stretching *τ*_str_. We express the time-averaged mean-field energy function for each cell in a form similar to the theory of Ref.[40], to compare the orientational behavior of the cells in a tissue to that of isolated cells. This is given by:

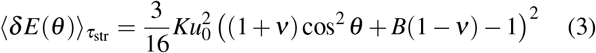

where *K* is an effective stiffness, and *B* is the anisotropy parameter of the cells defined in the SI Section 1. The anisotropy parameter *B* is a combination of the various elastic moduli along with the principal strain directions of the cells, and *B* → ∞ for isotropic cells, while *B* → 1 in the limit of strong anisotropy. The stability analysis using the energy function Eq.(3) suggests that there exist two regimes for the preferred direction *θ*\* (the minimum of the energy function) depending on the values of *B* and the Poisson ratio of the substrate 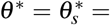 arccos 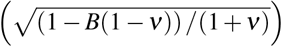 when *B* < 1/(1 − *ν*) and *θ*\* = *π*/2 when *B* ≥ 1/(1 − *ν*). We determine the value of the anisotropy parameter by mapping the geometrical parameters of the vertex model to the parameters of the elastic model. This yields

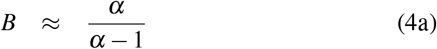

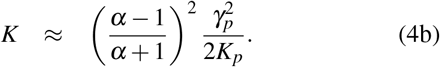

where *α* is the aspect ratio of the cells (see SI Section 1). Note that *K* would vanish for isotropic cells *α* = 1 for which the energy function becomes independent of the orientation angle. For our experiments, using the mean value *α* = 1.8, we estimate *B* = 2.25. Remarkably, the semi-empirical observation by analyzing a digitally stretched image (*B^e^* ≈ 2.29, see SI Section 1) validates this simple approach. For these parameters, we expect *θ*\* = *π*/2 in accordance with our experimental results. This behavior is different from the result obtained in [40] with isolated fibroblasts that displays a preferred angular orientation 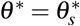 whose value depends on both *B* and the effective Poisson ratio at the center of the chip. Accordingly, in that work, the anisotropy parameter is fitted with a value *B* ≈ 1.1. Interestingly, Eq.(4) can also qualitatively reach this value for high aspect ratios that we would expect for fibroblasts even though we derived it using a vertex model. Similar to other studies with isolated cells having high aspect ratio *α* ≫ 1 for which *B* approaches unity report that the cells orient on average along the zero-strain direction [41–43] consistent with our framework. While this generality is remarkable, we should note that for confluent tissues with high *α*-value, our theory in Eq. (3) is no longer valid as one needs to consider steric interactions leading to long-range correlations between cells displaying nematic order.

As a whole, the effective dynamics of each cell can be written in the Langevin form:

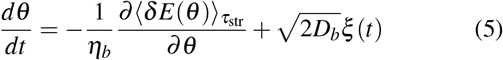

where *η_b_* is the basement membrane dependent friction coefficient and *ξ* (*t*) is a zero mean unit variance Gaussian white noise. The effective diffusive behavior with rotational diffusion constant *D_b_* originates from the binding rearrangements of the cells due to both intra-cellular processes and interactions with their surroundings. Similar approaches have been used in modeling cell migration polarity [44–46]. These dynamical equations can be transformed into a Fokker-Planck equation for the probability distribution *ρ*(*θ, t*):

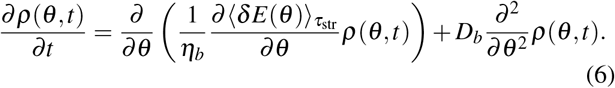

We can deduce the angular distribution of the cells as a function of time from the Fokker-Planck equation. To probe the tendency of orientation, we define an order parameter:

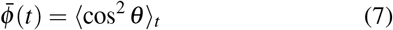

where the bracket 〈〉 *_t_* denotes an average performed over the distribution *ρ*(*θ, t*). For an ensemble with random orientations 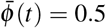, whereas 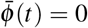 or 1 respectively for purely perpendicular and parallel orientations. Approximating the probability distribution *ρ*(*θ, t*) by a Gaussian function (see SI Section 1), we obtain a simple form of the order parameter as a function of time

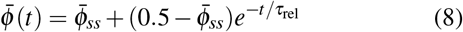

using the boundary conditions 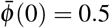 and 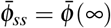 obtained from the experimental results. The relaxation time depends on both the stored elastic energy and the rotational diffusion constant, and is given by

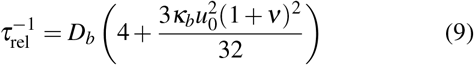

where *κ_b_ = K/η_b_D_b_*. For small deformations, the steady-state value of the order parameter becomes:

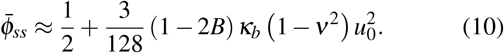

As our analysis shows *B* ≥ 1 (Eq.(4)) and since *ν* < 1, the second term is negative. Note that for isotropic cells as *α* = 1, *κ_b_* = 0 since *K* = 0 from Eq.(4) leading to 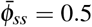 On the other hand, increasing *κ_b_* enhances the perpendicular orientation of cells as 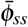 becomes lower than the random orientation value of 0.5.

In Fig. 3, we show the orientational response of the cells after 24 h of cyclic stretching (Fig. 3 B,C) and the time evolution of the order parameter (Fig. 3 E,F), together with the theoretical curve. During the reorientational dynamics, we observe a transition between a random distribution of orientations *ρ*(*θ,t*) and a Gaussian distribution centered around *θ* = *π*/2 as the total stretching time *t* increases. The order parameter 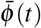 decreases with the stretching time until it converges to its steady-state (long-time) value 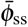 For the theoretical analysis, we write the rotational diffusion constant as *D_b_ = T_b_/η_b_* where *T_b_* is a coating-dependent effective temperature that stems from the dynamical activity of the cells channeled to the rotational degrees of freedom. Thus, *T_b_* can alter the amplitude of the angular fluctuations, independently from the rotational friction constant *η_b_*. This allows us to evaluate the coating-dependent changes of the steady-state angular distributions and of the relaxation time separately. The steady-state probability distribution has a Boltzmann form 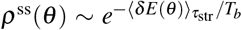 which is a function of *κ_b_* = *K/T_b_*. Setting *B* = 2.25, we obtain the best fit parameters as *κ*_cov_ = 128, *D*_cov_ = 0.053 h^−1^ for covalently coated, and *κ*_non-cov_ = 178, *D*_non-cov_ = 0.036 h^−1^ for non-covalently coated laminin basement membrane. Thus, we observe that the rotational diffusion coefficient in covalent condition is about 1.5 times higher than the non-covalent one. We may understand this behavior by the facilitated diffusion of cells as the surface density of laminin is higher in covalently bound laminin coating [33]. On the other hand, *κ_b_* has the inverse trend and the value *K/η_b_ = κ_b_D_b_* does not depend on the coating condition. This intriguing conservation allows us to make the following speculation. In our previous analysis, *K* is obtained from an effective mapping which depends on the Poisson ratio of the substrate *ν*, the geometrical parameters of the cells and the perimeter moduli *K_P_* and *γ_P_* (see SI Section 1). Since, we do not observe any difference in the morphologies of the cells in the covalent and non-covalent conditions, we can suppose that the values of *K_P_*, *γ_P_* and therefore of *K* are independent of the coating conditions, suggesting that the rotational friction constant *η_b_* is also coating-independent. Thus, ruling out the sensitivity of these parameters on coating conditions leads to *T*_cov_ ≈ 1.5*T*_non−cov_. This difference could be due to the fact that the energy transmission to rotational degrees of freedom is depending on the surface density which differs in the two coating conditions. Besides this hypothetical picture, we nevertheless conclude that the dynamical parameters remain within the same range for both coating conditions.

**Figure 3.**
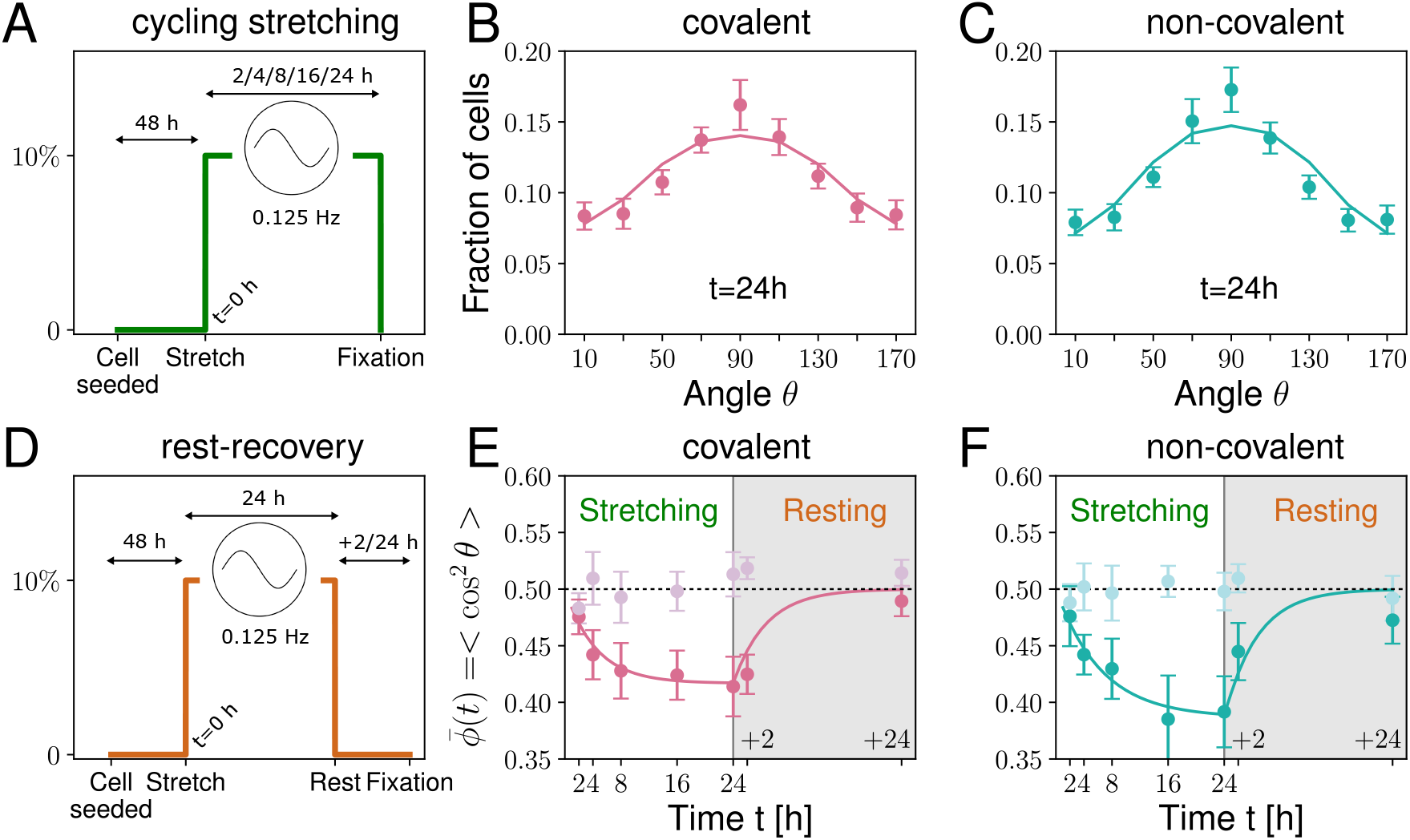
Orientational response of cells. (A) Time frame of cyclic stretching. The cyclic stretching parameters are displayed in Fig. 2. We show the angular distribution of cells after a total period of 24 h cyclic stretching (B) for covalent coating and (C) for non-covalent coating conditions. Dots with error bars are experimental results and the solid line is the theoretical curve (see fit parameters in text). (D) Time frame of rest-recovery experiments. We show the time evolution of the order parameter 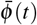 for different periods of cyclic stretching and rest-recovery periods (24 h stretching +2,+24 h rest) (E) for covalent coating and (F) for non-covalent coating conditions.

Finally, we performed rest-recovery experiments in which the confluent tissue first undergoes cyclic stretching during 24 hours and then left unstretched in position *u*_0_ = 0 for an additional *t* = 2 or 24 hours (see Fig. 3E). The evolution of the order parameter is shown in Fig. 3 E and F (gray regions) respectively for covalent and non-covalent conditions. We observe a recovery to the isotropic state with uniform angle distribution, showing that the angular response is reversible, in agreement with our theoretical description. We conclude that the dynamics is consistent with a rotational diffusion under an external mechanical stimulus. Thus, in the absence of potential, i.e., *u*_0_ = 0 in rest-recovery timeline, the diffusive relaxation rate 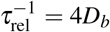 approximately captures the experimental trend for both conditions.

### The correlations are short-ranged and decay exponentially

We also studied the range of the spatial correlations of orientation to determine whether local fluctuations can have longrange effects. For two cells at a normalized distance *ℒ* ≡ *r/R* where *r* is the actual distance and *R* is the mean neighboring distance between neighboring cells in the tissue, we calculate the correlation function of the order parameter

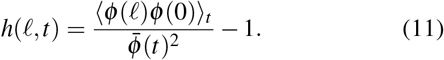

Here, the averaging is performed at a given time point over all cell pairs. We observe that the correlations in cell reorientation are not long-range. In Fig. 4 A, we give an example of the calculated correlation function in covalent conditions under a total of *t* = 24 h cyclic stretching. Moreover, the correlation function has an exponential decay *h*(*ℒ, t*) ∼ *e*^−l/*ξ*^ where the correlation length *ξ* appears independent of coating and stretching conditions (Fig. 4 B). In all cases, the correlation length is very short *ξ* ⪅ 1, which justifies our mean-field approach where we treated the dynamics of each cell independently.

On the other hand, the exponential decay of the correlations indicates that our system is away from algebraic ordering which is ubiquitous in two-dimensional systems with polar degrees of freedom where the correlations decay as a power law. In principle, the motility levels and the geometrical characteristics of Caco-2 cells can be the limiting factors. It would be interesting to see how these characteristics change in other cell types under cyclic stretching and whether a theory beyond the mean-field approach would be necessary.

**Figure 4.**
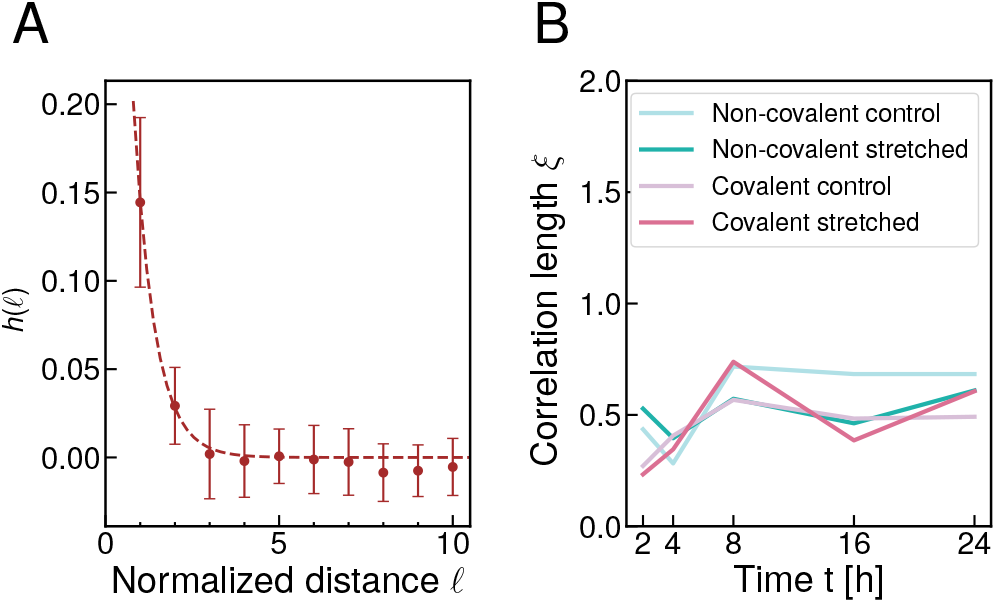
Range of orientation correlations. (A) Correlation function *h*(*ℒ*) as a function of normalized distance *ℒ* in the covalent coating condition after *t* = 24*h* of cyclic stretching. We observe that the correlations decay exponentially with a correlation length *ξ* ⪅ 1 and are even shorter range than the mean neighboring distance between cells. In (B), we show that *ξ* is independent of coating condition and total time of stretching.

## Conclusions and Outlook

In this work, we have studied the reorganization of intestinal tissue under cyclic stretching mimicking *in vivo* peristalsis. To do so, we designed an original stretcher along with specific PDMS chips compatible with conventional cell culture conditions allowing the application of a controlled cyclic mechanical deformation to an intestinal epithelial monolayer. Thus, we recapitulated the mechanical environment felt by epithelial cells in the intestine with a controlled stretching in terms of frequency, amplitude, duration and cyclic patterns. We have evaluated different laminin coating strategies (covalent and non-covalent) to understand their contribution to the signal mechano-transmission. Our results highlight a significant sensitivity to peristaltic-like stimuli of the angular orientation of the long axes of the cells with respect to the stretching axis with about 20% of the cells oriented with an angle ranging from 80 to 100°, whereas the other morphological parameters of the cells remain unchanged.

Earlier literature focused on the orientational response of isolated and highly contractile cells with high aspect ratio under cyclic stretching. The total yield and dynamics of cells are dependent on their elastic anisotropy, the stretching frequency, and the effective biaxiality ratio [40, 41]. Here, we have studied for the first time the orientational response of intestinal epithelium in details. In particular, peristaltic-like stimuli fall into the fast regime as it is faster than the topological rearrangement of cells. We show an equivalent approach from the vertex model which can be mapped to effective elastic theories. Thus, we propose a unifying theoretical framework to study the orientational dynamics for different types of cells by relating their geometrical properties to elastic ones. We model the orientational dynamics in tissue cells as a viscoelastic process along the rotational axis with a characteristic autocorrelation time *τ*_rel_ ∼ 5-7 hours. The discrepancy between covalent and non-covalent conditions is small, while the latter has slower relaxation dynamics. Thus, one can also argue the facilitated diffusion of cells on covalently coated laminin surface as there may be more available stable binding sites for cells [30], yet this would require microscopic examination of binding properties, which is beyond our scope in this paper. For our purposes, we conclude that the two conditions do not display a significant physiological difference.

Moreover, forces can also affect cell migration. Basson et al have studied the effect of repetitive deformation on wound closure of Caco-2 monolayer, showing that cell motility can be either stimulated or inhibited depending on the adhesion molecules nature [4]. The effect of stretching on cell migration has also been studied for many other cell types, such as Zhang et al who showed that cyclic stretching promotes bone marrow stem cell migration while slightly inhibiting cell invasion [47]. A recent work has shown that stretching forces could also induce the alignment of fibronectin secreted by human prostatic fibroblasts as well as a persistent co-directionality of cancer cells migration when co-cultured with stretched fibroblasts [48]. Taken together, these results suggest that stretching is prone to induce a change in cell migration properties. How this change of cell orientation under cyclic stretching contributes to an increased cell migration remains to be understood. Indeed, recent work from Krndija et al [1] has demonstrated that gut homeostasis relies on active migration of intestinal cells along the crypt to villus axis, opening fundamental questions about the gut homeostasis [49]. Among those, how biochemical and mechanical cues contribute to the polarity of the cell sheet is still under investigation. At this stage extrapolation between in vitro and *in vivo* studies remains elusive as the pattern of stretching forces experienced *in vivo* by intestinal cells is way more complex than cyclic stretching applied here. Besides this and as mentioned earlier, *in vivo* intestinal cells are grown on a 3D scaffold and are subjected to different curvatures being in a different configuration when compared with the flat PDMS membrane used here for cell culture. This is why the microfabrication process has been designed so that a stretchable 3D scaffold can easily replace the flat PDMS substrate. Our work paves the way for further studies performed with primary cells stretched on a 3D deformable scaffold to decipher the role of the mechanical cues on intestinal cell migration in 3D.

## Materials and Methods

### Chip fabrication and coating

A brass mold was prepared by micromilling (Minitech). The PDMS ship was molded directly on to the brass mold prepared with PDMS (Sylgard 184, Dow Corning) with a prepolymer /curing agent at ratio 10:1 (w/w). The mold has been designed to fabricate chips with a central reservoir for cell culture medium compatible with long term cells culture, a flat bottom membrane compatible with cell stretching (thickness 1mm) and connectors to be directly plugged the chips on the stretcher. Another possible refinement of the device relies on the possibility to further micro-mill the central part of the mold to change at will the 3D structure of the bottom of the chip. To perfectly control the brass mold positioning and consequently the chip wall thickness, all the parts of the molds were screwed and poured 7 mL of PDMS (1:10) and let to cure at 70°C for at least 3 hours. After demolding, chips were sterilized in an ethanol bath (70%). Then we use two different coating protocols :

i. **non-covalent coating** requires first the activation of the surface by a *O*_2_ plasma treatment, then a solution of laminin (Natural Mouse, from Invitrogen 23017-015) at 0.02 mg/mL is incubated for 1 hour at 37°C and then washed 3 times with water. In this condition, the laminin coating mainly relies on electrostatic, hydrogen and Van der Waals interactions.
ii. **covalent coating** requires first the activation of the surface by a *O*_2_ plasma treatment, then a solution of (3-aminopropyl) triethoxy-silane (APTES) at 0.5% (v/V) is incubated for 30 minutes, then rinsed 3 times the with deionized water, and the chip incubated for 30 minutes with a solution of cross-linker glutaraldehyde at 2% (v/v). Finally, the chip is rinsed overnight with deionized water, sterilized with ethanol and incubated with laminin (0.02 mg/mL) for 1 hour at 37°C.

### Cell culture

Caco-2 cells were plated on the laminin coated chips with a cell concentration of 200 000 cells/chip, let adhere to the surface for 24 hours then medium is refreshed every 24 hours. After 48 hours of culture, cells reach confluency and three chips are placed on the stretcher and three are let free as control conditions, all were placed in a conventional cell culture incubator. For all the experiments the medium used was DMEM (1*X*) + GlutaMAX (ThermoFisher scientific 31966-021) with 10% of Fetal Bovine Serum (ThermoFisher scientific 26140079), 1% of Antibiotico-antimycosique (ThermoFisher scientific 15240062) and 1% of MEM Non-Essential Amino Acids Solution (ThermoFisher scientific 11140035) [50].

### Staining protocol

We fix the cells using a solution of PFA/PBS at 4% (v/v) for 30 minutes at room temperature. Then we rinse 3 times with PBS during 30 minutes and keep the chip in PBS overnight. We incubate the chips with Triton X-100/PBS at 0.5% (v/v) for 5 min at room temperature (500uL of permeabilization solution per chip). We incubate the chips with the *blocking buffer* (0.2% of Triton X-100/2% of BSA/3% of normal goat serum) for 2h at room temperature (500uL Blocking buffer per chip). We incubate the chips with primary antibodies in the *working buffer* (0.2% of Triton X-100*/*0.1% of BSA/0.3% of normal goat serum) for 2h at room temperature. The primary antibody ZO1 monoclonal (ThermoFisher scientific 33 – 9100) was used at 1/75th dilution and 300uL of solution per chip were used. We wash 3 times with PBS for 30 min each at room temperature. We incubate the chips with secondary antibodies in the *working buffer* for 1h at room temperature. The secondary antibody Goat anti-Mouse Alexa Fluor Plus 647 (ThermoFisher scientific *A*32728) was used at 1/100th dilution and 300 uL of solution per chip were used. We wash 4 times with PBS for 30 min each at room temperature. We mount the samples using a mounting medium (ThermoFisher scientific *P*36931). Then we store the slides horizontally at 4 °C until image acquisition.

### Image acquisition and analysis

After staining, images from 5 different fields per chip were taken using a 10× objective, on a confocal microscope. All experiments were repeated twice. Therefore, 30 images were obtained for every experiment for the stretched and control conditions. The 10× objective induces a wide field of view (1163.64 μm × 1163.64 μm). The graphics in the main text represent the distribution or the mean value inside each image and the error bars are the standard deviation between those 30 images. These measures were obtained using ImageJ software [51], and in particular the plugin Tissue Analyzer [52].

## Acknowledgments

We acknowledge support from the Labex CelTisPhyBio (ANR-11-LABX-0038, ANR-10-IDEX-0001-02) and we also acknowledge DIM ELICIT for funding. This work has received the support of “Institut Pierre-Gilles de Gennes” (laboratoire d’excellence, “Investissements d’avenir” program ANR-10-IDEX-0001-02 PSL, ANR-10-LABX-31, ANR-10-EQPX-34, ANR project HOMEOGUT and Holifab (H2020-EU.2.1.2. H2020-NMBP-PILOTS-2017, grant agreement 760927).

## Author contributions

L.G., E.I., J.-L. V., D.M.V., J.-F.J., S.D. participated to the design research design; L.G. build the experimental platform; C.C. characterise the experimental platform; L.G.,M.B.-D. performed experiments ; L.G., and E.I participated to data analysis and interpretation, E.I. and J.-F.J. developed the theoretical models; and L.G, E.I., D.M.V., J.-F.J., S.D. wrote the paper.

