## Supplementary Information for "Evolution of a confluent gut epithelium under cyclic stretching"

### Supplemental Material for "Evolution of a confluent gut epithelium under cyclic stretching"

#### S1. MODELING THE ORIENTATIONAL RESPONSE OF EPITHELIAL CELLS UNDER STRETCHING

We follow the assumptions stated in the main text. These suggest: (i) Cells deform together with the substrate and hence passively follow the stretch direction at short times ($t \sim \tau_{\rm str}$) where $\tau_{\rm str} = 2\pi/\omega$ is the period of stretching. (ii) At long timescales ($t \gg \tau_{\rm str}$), the cells undergo a rotational diffusion. Thus, the cells release this energy through a dissipative process in which they rearrange their shapes and boundaries. Under these assumptions, we need to calculate the mechanical energy stored by the cells due to deformation of the substrate. We first use a vertex model. Then, we show a mapping to an effective continuum elastic model similar to that for isolated cells such as fibroblasts. We highlight the similarities and the differences between isolated cells and cells in a tissue.

##### A. Mechanical energy

###### *1. Vertex model and mean energy injected per cell*

We first investigate the cell deformations through the vertex model which is widely used for studying the dynamics of 2-dimensional confluent tissues. For a tissue containing $N$ cells, the total mechanical energy is given by $E = E(\{\mathbf{r}_i\})$[1, 2],

$$E_{\rm tot} = \sum_{i=1}^{N} K_A \left(A_i - A_0\right)^2 + \gamma_P P_i + K_P P_i^2 \tag{S1}$$

where each cell is labeled by an index $i$ while $A_i$ and $P_i$ are respectively the area and the perimeter of cell $i$. The energy related to changes in area is quadratic around an optimal cell area $A_0$ fixed by the interplay between cell volume incompressibility and the limiting resistance to height fluctuations. The second term proportional to the cell perimeter $P_i$, is a line tension due to both cell-cell adhesion and cortical tension. We can separate these two contributions to the line tension as $\gamma_P = -\gamma_P^A + \gamma_P^T$ : if $\gamma_P < 0$, then adhesion dominates and vice versa. The last term proportional to $K_P$ is the elastic contribution due to acto-myosin contractility, and is quadratic in the perimeter of the cell $P_i$. Thus, the cells have a preferred perimeter $P_0 = -\gamma_p/2K_p$ which we consider to be identical for each cell with the choice of homogeneous $K_P$ and $\gamma_P$ values. The ground state configuration of the tissue is found by minimizing the total mechanical energy $E_{\rm tot}$. Here, we focus on the change of total energy function due to the external mechanical strain applied on the substrate. We can express this perturbation for each cell as a function of cell's long axis orientation angle. Upon stretching, the area of the cell $A_i$ varies as $A_i \to A_i(1 + u(t))(1 - \nu u(t))$ and hence it is independent of the orientation. By contrast, the perimeter varies as a function of orientation angle and the change in total energy is given by:

$$\delta E_{\rm tot}(t) = \sum_{i=1}^{N} K_p P_i^2 \left(p_i(\theta_i, u(t))^2 - 1\right) + \gamma_P P_i \left(p_i(\theta_i, u(t)) - 1\right) + \text{angle-indep. terms} \tag{S2}$$

where $P_i(\theta_i, u(t))$ is the perimeter of $i^{\rm th}$ cell with an orientation $\theta_i$ under strain $u(t)$, and $p_i(\theta_i, u(t)) = P_i(\theta_i, u(t))/P_i$ is its ratio in reference to the perimeter value $P_i$ under no strain. Therefore, the minimization of $\delta E_{\rm tot}(t)$ corresponds to a minimization of the change of the cell perimeter upon stretching.

In order to obtain the mechanical energy injected per cell during stretching, we use a mean-field approach. Accordingly, we set $P_i = P_0 = -2\gamma_P/K_P$ and obtain the average energy per cell from Eq.(S2)

$$\delta E(\theta, t) = \frac{\gamma_P^2}{4K_p}(p(\theta, u(t)) - 1)^2. \tag{S3}$$

Thus, $\delta E(\theta, t)$ is a function of the cell orientation angle and we drop the index $i$ considering the dynamics of each cell identical yet dependent on $\theta$.

##### 2. *Effective elastic theory*

We next consider the cells as anisotropic elastic materials along the lines of the model introduced in Ref. [3] that we reformulate in a slightly different form using Cartesian coordinates. In continuum elasticity, the stored energy due to the deformation of an elastic material is given by [4]:

$$\mathcal{H} = \frac{1}{2}\tilde{\lambda}_{ijkl} u_{ij} u_{kl} \tag{S4}$$

where $\tilde{\lambda}_{ijkl}$ is the elastic modulus tensor, $u_{ij}, u_{kl}$ is the strain or deformation tensor. The indices $i, j, k, l$ denote the Cartesian components and we use the Einstein summation convention. We use the notation $(x^{\rm lab}, y^{\rm lab})$ for the coordinates in the lab reference frame and $(x, y)$ for the coordinates in the reference frame of the cell as shown in Fig.S1. Using the symmetries for plane-stress, we obtain the elastic strain energy of the cell that is integrated over the area of the cell:

$$\mathcal{H}_{cell} = \frac{1}{2}\lambda_{xxxx} u_{xx}^2 + \frac{1}{2}\lambda_{yyyy} u_{yy}^2 + \lambda_{xxyy} u_{xx} u_{yy} + 2\lambda_{xyxy} u_{xy}^2 \tag{S5}$$

which involves 4 independent moduli associated to the anisotropic response of the cells, and $\lambda_{ijkl} = A_0 \tilde{\lambda}_{ijkl}$ has units of energy. Stretching deforms each point on the substrate by a time varying strain along $x^{\rm lab}$-direction with frequency $\omega$: $u_{xx}^{\rm lab}(t) \equiv u(t) = \frac{1}{2}u_0(1 - \cos\omega t)$. The resulting compression along the $y^{\rm lab}$-axis is $u_{yy}^{\rm lab}(t) = -\nu u(t)$ where $\nu$ is the Poisson ratio of the substrate. More generally, if one controls the elastic strains along $x^{\rm lab}$ and $y^{\rm lab}$ independently, $\nu$ should be replaced by the measured magnitude of biaxiality strain ratio $u_{yy}^{\rm lab}/u_{xx}^{\rm lab}$ at the center of the chip. The elastic strains due to the stretching of the substrate can be transformed to the cell's reference of frame as

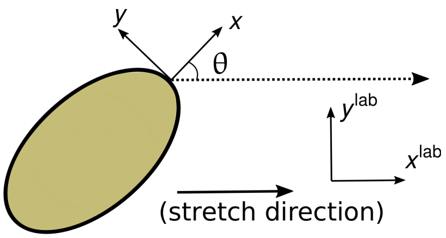


FIG. S1. Illustration of the coordinate axes. In our analysis, each cell is described by a best-fit ellipse with semi-major axis $a$ (along $x$-direction), semi-minor axis $b$ (along $y$-direction), and angle of orientation $\theta$ which is the angle between the semi-major axis and the stretching direction. $(x^{\mathrm{lab}}, y^{\mathrm{lab}})$ corresponds to lab frame, where the stretching occurs along $x^{\mathrm{lab}}$-direction. The cell's principal axes frame $(x, y)$ can be obtained with by rotation of the lab frame by an angle $\theta$ which is defined so that $x$ is the long-axis of the cell.

$u_{xx}(t) = u(t)\left(\cos^2\theta - \nu\sin^2\theta\right)$, $u_{yy}(t) = u(t)\left(\sin^2\theta - \nu\cos^2\theta\right)$ and $u_{xy}(t) = u(t)\,(1+\nu)\cos\theta\sin\theta$. Inserting these strains, we obtain the angle-dependent part of the elastic energy as:

$$\mathcal{H}_{cell}(\theta, t) = \frac{1}{2}Ku^2(t)\Big((1+\nu)\cos^2\theta + B(1-\nu) - 1\Big)^2. \tag{S6}$$

Thus, the stored elastic energy per cell depends on only two cell-related parameters. The anisotropy parameter $B$ and the elastic coupling constant $K$ are given by:

$$B = \frac{\lambda_{xxxx} - (\lambda_{xxyy} + 2\lambda_{xyxy})}{\lambda_{xxxx} + \lambda_{yyyy} - 2(\lambda_{xxyy} + 2\lambda_{xyxy})} \tag{S7a}$$

$$K = B^{-1}\left(\lambda_{xxxx} - (\lambda_{xxyy} + 2\lambda_{xyxy})\right). \tag{S7b}$$

The anisotropy parameter $B$ is dimensionless and the modulus $K$ has units of energy. The energy function Eq.(S6) has the same form obtained in Ref.[3] for isolated cells (note that the definition of $K$ is different).

##### 3. *Crossover between preferred orientations determined by elastic anisotropy*

In energy function given in Eq.(S6), $B$ is the key parameter that determines the preferred direction for long-axis orientation for cells. $B$ itself is determined by the level of elastic anisotropy; weakly anisotropic cells reach $B \gg 1$ and the energy function becomes independent of the orientation angle, whereas strong anisotropy leads to $B \to 1$ which we will illustrate further below. The stability analysis of the elastic energy Eq.(S6) shows that there are two regimes for the preferred cell orientation (given by the minimum of the elastic energy):

$$\theta^* = \begin{cases} \theta^*_{\mathrm{s}} = \arccos\left(\sqrt{\frac{1-B(1-\nu)}{1+\nu}}\right) & \text{when } B < 1/(1-\nu), \\ \pi/2 & \text{when } B \geq 1/(1-\nu). \end{cases} \tag{S8}$$

Thus, when $B < 1/(1-\nu)$, the preferred angle $\theta^*_{\mathrm{s}}$ depends on both the substrate (PDMS) Poisson ratio $\nu$ and the cell anisotropy parameter $B$. For strongly anisotropic cells, $B \to 1$ and hence

$\theta_s^*$ approaches the zero-strain orientation, i.e., $\theta_s^* \to \arccos\left(\sqrt{\frac{\nu}{1+\nu}}\right)$. The isolated fibroblast cells orient in mirror-image directions given by a value of $\theta^*$ that depends on both $\nu$ and $B$ which allows to determine the value of $B$ from experiments [3]. By contrast, for cells in a confluent tissue, the anisotropy is weaker and our results show that the cells tend to orient perpendicular to the stretch direction. Thus, the observation of the preferred angle is not a determinant to deduce $B$ except that we should expect $B \geq 1/(1-\nu)$. The rheology experiments on the tissue can be used to measure the elastic moduli $\lambda_{ijkl}$ in order to obtain $B$ and $K$, however, they would yield tissue level properties. On the other hand, we are interested to distinguish the anisotropy at the cell level which is the main feature shaping the orientational response of cells. In the following, we estimate the anisotropy parameter $B$ purely from the geometrical properties of tissue cells by using the vertex model.

##### 4. *Determination of the anisotropy parameter B from the vertex model*

We can map the mean energy per cell obtained from the vertex model in Eq.(S3) to the form of energy function in Eq. (S6) and obtain the anisotropy parameter $B$. In order to do so, we need to estimate the fractional change in perimeter as a function of orientation angle $\theta$ and applied strain $u(t)$.

*a. Simple ellipse assumption:* The simplest approach is to approximate each cell by an ellipse with short and long axes $a$ and $b$ respectively. If we then approximate the perimeter by $P \approx \pi(a+b)$ and apply strains along the axes of the cell, we obtain $\delta E(\theta, t) = \mathcal{H}_{cell}(\theta, t)$ with parameters

$$B = \frac{\alpha}{\alpha - 1} \tag{S9a}$$

$$K = \left(\frac{\alpha - 1}{\alpha + 1}\right)^2 \frac{\gamma_p^2}{2K_p}. \tag{S9b}$$

where we define the aspect ratio $\alpha = a/b$. Our result remarkably agrees with isolated cell results so that $B \to 1$ for strong anisotropy as $\alpha \gg 1$ whereas $K$ vanishes for isotropic systems $\alpha = 1$ and the energy function becomes independent of the orientation angle. For our experimental system, using the observation that the mean value of the aspect ratio is $\alpha = 1.8$, we obtain $B = 2.25$.

*b. Semi-empirical approach:* We also measured the value of the anisotropy parameter $B$ using empirical considerations. From our image analysis, we first obtain the perimeter values $P_i(\theta_i)$ and orientation angle $\theta_i$ for each cell. Then, we stretch the same image digitally with $u_{xx}^{\text{lab}} = u$, $u_{yy}^{\text{lab}} = -\nu u$ and perform an image analysis in order to obtain the stretched perimeter value $P_i(\theta_i, u)$ for each cell. Then we define $\delta p(\theta_i, u) = \left(P_i(\theta_i, u) - P_i\right)/P_i$, and perform a chi-square fitting to a function,

$$\delta p(\theta_i, u) = c_1 \cos^2 \theta_i + c_2. \tag{S10}$$

using $u = 0.1$ with fitting coefficients $c_1$ and $c_2$. Now, if we call the empirical anisotropy parameter $B^e$ and use the quadratic form in Eq.(S6), we find the following relation

$$\frac{c_1}{c_2} = \frac{2(1+\nu)}{B^e(1-\nu-1/B^e)}. \tag{S11}$$

from which we obtain $B^e \approx 2.29$.

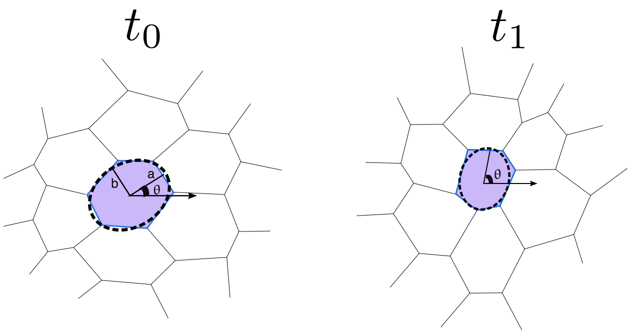


FIG. S2. Illustration of rotational diffusion of the long axis of a cell. Even in the absence of mechanical strain, tissue cells exhibit shape fluctuations by rearranging their bonds and hence their shape parameters are different at times $t_0$ and $t_1$. Accordingly, we model the fluctuations in orientation of the long-axis as a rotational diffusion.

##### B. Rotational Dynamics

The configuration of the cells in the tissue constantly deviates from the ground state of the entire network due to active processes [5]. The system displays many alike preferred configurations. As illustrated in Fig.S2, we link this behavior including all the topological transitions (T1 and T2 transitions [6], and cell divisions) to the rotational diffusion of the long axis of cells. If we now define a basement-membrane-dependent rotational diffusion coefficient $D_b$ and friction constant $\eta_b$, we obtain the Langevin dynamics:

$$\frac{d\theta}{dt} = -\frac{1}{\eta_b}\frac{\delta E(\theta,t)}{\partial\theta} + \sqrt{2D_b}\xi(t) \tag{S12}$$

where $\xi(t)$ is a zero mean unit variance Gaussian white noise. From the Langevin equation, we derive a Fokker-Planck equation to study the time evolution in terms of the probability distribution function $\rho(\theta,t)$ of the orientations. From the Fokker-Planck equation, we study in details the time evolution of the tissue in the two periods of motion: i) cells moving under periodic stretching and ii) resting period which follows the stretching period and which the cell orientation relaxes in the absence of stretch. Having estimated the elastic anisotropy $B = 2.25$, we are left with 3 fitting parameters $\{\eta_b, D_b, K\}$ to determine.

i) *Periodic stretching:* We observe that rotational diffusion is a much slower process than the stretching frequency, $D_b \ll \pi^2\omega$. This allows us to average over one period $\tau_{\rm str}$ the time varying strain $u^2(t)$. We thus replace $u^2(t)$ by its time average $\langle u^2(t)\rangle = \tau_{\rm str}^{-1}\int_0^{\tau_{\rm str}} u^2(t)dt = (3/8)u_0^2$ in Eq.

(S6). As a result, we obtain the time evolution of the probability distribution of cell orientations $\rho(\theta,t)$ as:

$$\frac{\partial\rho(\theta,t)}{\partial t} = -\frac{3K}{8\eta_b}u_0^2(1+\nu)\frac{\partial}{\partial\theta}\sin 2\theta\Big((1+\nu)\cos^2\theta + B(1-\nu) - 1\Big)\rho(\theta,t) + D_b\frac{\partial^2}{\partial\theta^2}\rho(\theta,t). \quad \text{(S13)}$$

This Fokker-Planck equation can be solved numerically starting from a uniform distribution at $t=0$, i.e., $\rho(\theta(0)) = 1/\pi$ with periodic boundary conditions such that $\rho(0,t) = \rho(\pi,t)$ and $\rho_\theta(0,t) = \rho_\theta(\pi,t)$ where $\rho_\theta = \partial\rho/\partial\theta$. We focus on the time evolution of the order parameter $\bar{\phi}(t) = \langle\cos^2\theta\rangle_t$ where the average is taken over the probability distribution $\rho(\theta,t)$. As mentioned in the main text, the order parameter measures the average orientation of the cell ensemble in the tissue; $\bar{\phi}(t) = 0.5$ for random orientations, $\bar{\phi}(t) = 1$ for purely parallel orientations and $\bar{\phi}(t) = 1$ for purely perpendicular orientations. It can be obtained using Eq.(S13) by multiplying both sides with $\cos^2\theta$ and integrating over the angle $\theta$. This yields

$$\frac{d}{dt}\langle\cos^2\theta\rangle_t = -k_0 - k_1\langle\cos^2\theta\rangle_t - k_2\langle\cos^4\theta\rangle_t - k_3\langle\cos^6\theta\rangle_t \quad \text{(S14)}$$

where $k_0, k_1, k_2, k_3$ are the following constants:

$$k_0 = -2D_b, \quad \text{(S15a)}$$

$$k_1 = 4D_b\left(1 + \frac{3}{8}\kappa u_0^2(1+\nu)\left(B(1-\nu)-1\right)\right), \quad \text{(S15b)}$$

$$k_2 = \frac{3}{2}D_b\kappa u_0^2(1+\nu)\left(2+\nu-B\left(1-\nu\right)\right), \quad \text{(S15c)}$$

$$k_3 = -\frac{3}{2}D_b\kappa u_0^2\left(1+\nu\right)^2, \quad \text{(S15d)}$$

where

$$\kappa_b = K/\eta_b D_b. \quad \text{(S16)}$$

Approximating $\rho(\theta,t)$ by a Gaussian distribution centered around $\pi/2$, we can express the averages of higher moments in terms of the ensemble average of the order parameter at time $t$ as $\bar{\phi}(t) = \langle\phi\rangle_t$. To do so, we first express:

$$\cos^{2m}\theta = \frac{1}{2^{2m-1}}\left[\frac{1}{2}\binom{2m}{m} + \sum_{k=0}^{m-1}\binom{2m}{k}\cos\left[2(m-k)\theta\right]\right] \quad \text{(S17)}$$

where $\binom{x}{y}$ denotes the binomial coefficients. Then, we use the fact that $\langle\cos 2m\theta\rangle_t \sim (\langle\cos 2\theta\rangle_t)^{m^2}$ for a Gaussian distribution. As $\langle\cos 2\theta\rangle_t \ll 1$, we ignore higher order contributions that have $m > 1$, and obtain:

$$\bar{\phi}(t) = \frac{\langle\cos 2\theta\rangle_t + 1}{2}, \quad \text{(S18)}$$

$$\langle\cos^4\theta\rangle_t \approx \bar{\phi}(t) - \frac{1}{8}, \quad \text{(S19)}$$

$$\langle\cos^6\theta\rangle_t \approx \frac{15}{16}\bar{\phi}(t) - \frac{5}{32}. \quad \text{(S20)}$$

We may then rewrite Eq.(S14) as

$$d\bar{\phi}(t)/dt = -\tau_{\rm rel}^{-1}\left(\bar{\phi}_t - \bar{\phi}_{ss}\right) \tag{S21}$$

where $\bar{\phi}_{ss} = \tau_{\rm rel}\left(\frac{1}{8}k_2 - k_0 + \frac{5}{32}k_3\right)$ is the steady-state value of the order parameter and $\tau_{\rm rel}^{-1} = k_1 + k_2 + \frac{15}{16}k_3$ is the relaxation rate. Using Eqs.(S15), we obtain

$$\tau_{\rm rel}^{-1} = D_b\left(4 + \frac{3}{32}\kappa_b u_0^2(1+\nu)^2\right) \tag{S22}$$

whereas the steady-state value of the order parameter for small deformations ($\kappa u_0^2 \ll 128/3$) becomes,

$$\bar{\phi}_{ss} \approx \frac{1}{2} + \frac{3}{128}\left(1-2B\right)\kappa_b\left(1-\nu^2\right)u_0^2. \tag{S23}$$

The solution of the differential equation for the order parameter is

$$\bar{\phi}(t) = \bar{\phi}_{ss} + (0.5 - \bar{\phi}_{ss})e^{-t/\tau_{\rm rel}} \tag{S24}$$

using the boundary conditions $\bar{\phi}(0) = 0.5$ and $\bar{\phi}(\infty) = \bar{\phi}_{\rm ss}$. With this form, we can determine the effective relaxation time $\tau_{\rm rel}$ by direct fit of the experimental data.

ii) *Resting:* During the resting period ($t$=24-48 h in Fig. 2 main text), the cell orientation relaxes in the absence of any external potential. Thus, $\delta E = 0$ ($u_0 = 0$) and the initial distribution is $\rho(\theta, t = 24\text{ h}) = \rho_{ss}(\theta)$. This is a pure rotational diffusion process,

$$\frac{\partial\rho(\theta,t)}{\partial t} = D_b\frac{\partial^2}{\partial\theta^2}\rho(\theta,t). \tag{S25}$$

Then, by applying the same manipulations as above, we have the form Eq.(S14), with $k_2 = k_3 = 0$ and $k_1 = 4D_b$, whose solution reads:

$$\bar{\phi}^{\rm rest}(t) = 0.5 + (\bar{\phi}_{ss} - 0.5)e^{-4D_b t} \tag{S26}$$

At long times, we recover $\bar{\phi}_t^{\rm rest} = 0.5$, i.e., a random cell orientation. In Fig.2 (main text) we observe a good agreement of this dynamics with the experimental results.

#### S2. STATIC STRETCHING EXPERIMENTS

To probe the behavior of a Caco-2 cell monolayer in response to static stretching, we use the following procedure: first, the intestinal cells are seeded on the chips 48 hours prior the beginning of the stretching experiments. After 48 hours of culture, the cells form a confluent monolayer, that we refer to as the reference state (RS). We study here properties of the whole epithelium rather than the isolated cell's response to stretch. The chips are then subject to a static elongation in a pulled state (PS) for 4 to 24 hours. For sustained elongation, the dynamics of epithelial cells take place during this stage PS. After each selected time period (4h, 24h), the stretcher is brought back to the reference state and the cells are fixed. This final stage where the cells are fixed is referred to as the quenched state (Q) at which we perform the measurements.

These experimental results confirm our assumption on the existence of two different time regimes where the cells have different behaviors: i) Cells remain attached and hence passively follow the stretch direction at short times. In principle, the stretcher device is moved from RS to PS and vice versa (PS to Q) in sufficiently short time that we observe only a passive behavior of the cells, i.e., the cells follow the substrate deformation. Thus, by applying static strains $u_{xx}$ and $u_{yy}$ on each cell there is one to one mapping from Q to PS setup and vice versa. ii) At long timescales, cells undergo a rotational diffusion and recover an isotropic distribution corresponding to random orientations. Once the cells start detaching, the effect of the mechanical stimulus is not maintained, and hence no mechanical energy is stored. In other words, the set point for zero strain becomes PS. Therefore, $\delta E(\theta, t) = 0$ and using Eq.(S12), we have the Langevin form:

$$\frac{d\theta}{dt} = \sqrt{2D_b}\xi(t) \tag{S27}$$

where $D_b$ is the effective rotational diffusion constant of a cell in the confluent tissue and $\xi(t)$ is a zero mean unit variance Gaussian white noise.

Using the first assumption, we expect cells to orient parallel to the stretching axis at initial times, while in the absence of mechanical energy, they reorient in random directions at long times in LS following Eq.(S27). This is shown in detail on Fig. S3 in terms of fraction of cells as a function of orientation angle $\theta$. As shown in Fig. S3 F, the initial stretching leads to parallel orientation $\bar{\phi}^{\mathrm{PS}}(0) > 0.5$, while the orientational dynamics which is on the order of hours follows:

$$\bar{\phi}^{\mathrm{PS}}(t) = 0.5 + 0.07e^{-4D_b t} \tag{S28}$$

where $D_b$ is the effective rotational diffusion constant.


[1] L. Hufnagel, A. A. Teleman, H. Rouault, S. M. Cohen, and B. I. Shraiman, On the mechanism of wing size determination in fly development, Proceedings of the National Academy of Sciences **104**, 3835 (2007).

[2] R. Farhadifar, J.-C. Röper, B. Aigouy, S. Eaton, and F. Jülicher, The influence of cell mechanics, cell-cell interactions, and proliferation on epithelial packing, Curr. Biol. **17**, 2095 (2007).

[3] A. Livne, E. Bouchbinder, and B. Geiger, Cell reorientation under cyclic stretching, Nature Communications **5**, 3938 (2014).

[4] L. D. Landau, A. Kosevich, L. P. Pitaevskii, and E. M. Lifshitz, *Theory of elasticity* (Butterworth, 1986).

[5] D. B. Staple, R. Farhadifar, J.-C. Röper, B. Aigouy, S. Eaton, and F. Jülicher, Mechanics and remodelling of cell packings in epithelia, The European Physical Journal E **33**, 117 (2010).

[6] D. Weaire and N. Rivier, Soap, cells and statistics—random patterns in two dimensions, Contemporary Physics **25**, 59 (1984).

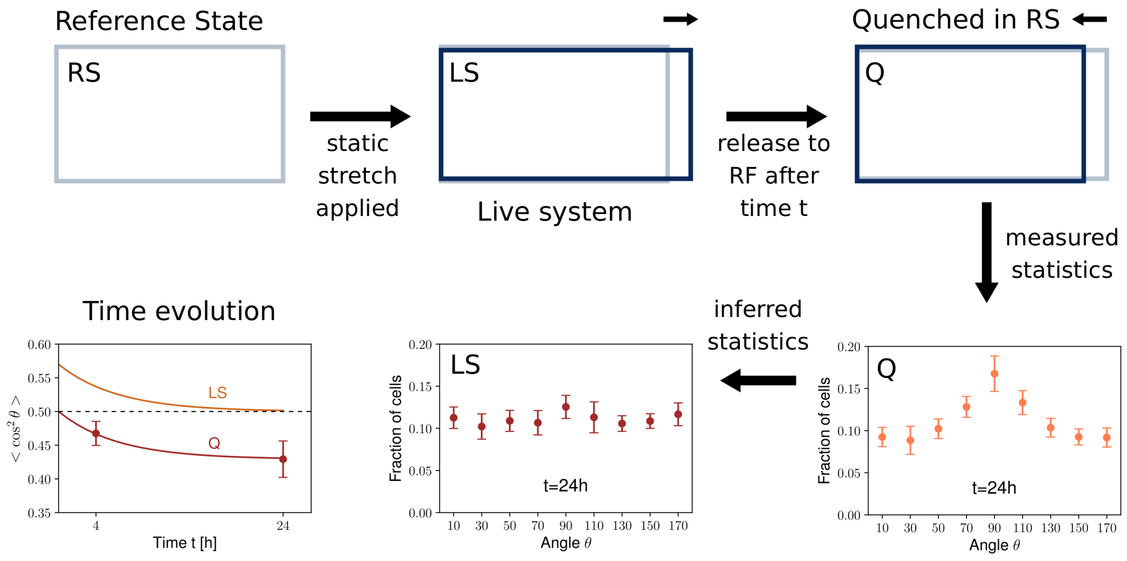


FIG. S3. **Static stretching experiments.** Cells are prepared on the regular chip which we name as the reference state (RS). Static stretch is applied while the cells are left to their dynamics for $t$ hours (Live system:LS). After $t$ hour, we release the applied stress and the cells are fixed right after. Thus, the system is quenched in RS, which we refer as Q. The statistics are obtained in Q, and then by applying the (reverse) strain (purely geometrical), we obtain the inferred statistics. In leftmost bottom graph, we show the time evolution of the order parameter $\bar{\phi}(t) = \langle\cos^2\theta\rangle_t$ in both frames.
